# Tear secretion *via* a paracellular pathway in lacrimal gland is regulated by myosin-mediated modulation of tight junction permeability

**DOI:** 10.1101/2024.11.06.622354

**Authors:** Yuta Ohno, Gaizun Hu, Dominik Robak, W. Sharon Zheng, Seham Ebrahim

## Abstract

“Dry eye”, characterized by symptoms of ocular discomfort and visual disturbances due to decreased tear secretion, affects 16 million Americans. Yet, there is currently no cure for dry eye as the mechanistic details of water secretion in the tear-producing lacrimal gland have not been fully elucidated. While a transcellular water secretion pathway *via* water channels like AQP5 has been reported, the existence and function of a paracellular pathway *via* tight junctions between epithelial cells remains controversial. The actomyosin cytoskeleton localizes to the apical junctions of epithelial cells across organs and regulates tight junction integrity. Here, we report that non-muscle myosin IIC (NMIIC) is enriched at apical junctions of ductal epithelial cells in the lacrimal gland, leading us to hypothesize that NMIIC regulates tear secretion through modulation of tight junction permeability. Consistent with this hypothesis, we found that tear volume after carbachol stimulation was significantly increased in mice lacking NMIIC, and levels of the tight junction protein ZO-1 were significantly reduced. Furthermore, pharmacological activation of NMIIC by 4-Hydroxyacetophenone in wildtype mice significantly inhibited tear secretion. In summary, our findings reveal a paracellular water secretion pathway in the lacrimal gland, which is regulated by NMIIC-mediated modulation of ductal cell tight junctional permeability, and can be targeted by small molecules.

**Significance Statement:** While dry eye affects more than 16 million Americans, there is currently no cure as the mechanisms underlying tear secretion are incompletely understood. Here, we report first evidence for the existence and function of a paracellular water pathway, in which water flows between cells, in the lacrimal gland. We also show that this pathway is directly regulated by the modulation of tight junction permeability by non-muscle myosin IIC (NMIIC). This study thus identifies a new mechanism for exocrine secretion, which can be targeted towards developing treatments for dry eye/mouth syndrome.

## Introduction

Tears play a critical role in corneal health by supplying oxygen and nutrients, and protecting from bacteria. Reduced tear secretion can lead to “dry eye”, causing blurred vision, burning or itching in the eye and sensitivity to light (1). Dry eye leads to reduced quality of life in approximately 16 million Americans (2). The causes of reduced tear secretion include Sjögren’s syndrome (3), the use of drugs having anticholinergic effects (4), radiotherapy for head-and-neck tumors (5), metabolic syndrome (6) and prolonged exposure to video display terminals (7). There is currently no causal cure for dry eye due to the incomplete elucidation of the mechanism of tear secretion. Tears are produced in the lacrimal gland, which mainly comprises two different cell-types: acinar cells, which contain secretary granules, and ductal cells that form a tubular network along which secretions are delivered to the surface of the eyeball. There are currently two putative water secretion pathways that have been proposed to exist: a transcellular water pathway *via* water channels such as AQP5, and a paracellular water pathway *via* tight junctions. In the lacrimal gland, a half reduction of tear secretion in AQP5 deficient mice (8, 9) provided evidence for the transcellular pathway, but also implies the involvement of other pathways, as tear secretion was not completely abolished. However, the existence and function of the paracellular water pathway remains controversial.

Myosin is a superfamily of motor proteins that hydrolyze ATP to generate mechanical forces and regulate actin filament dynamics and architecture. The myosin II subfamily of conventional myosins includes skeletal, cardiac and smooth muscle myosins, and non-muscle myosin-II (NMII) paralogs (10). NMII, ubiquitously expressed in non-muscle cells, has several roles in regulating cell movement, division, morphology (11). NMII has recently also been reported to play a critical role in tight junction formation in zebrafish embryos (12). There are three NMII paralogs in mammalian cells; NMIIA, NMIIB, and NMIIC which are derived from three different genes, MYH9, MYH10, and MYH14, respectively (13). NMIIA and NMIIB are reported to connect ZO-1 *via* cell junctional proteins *in vitro* (14), and distribution of junctional NMIIB was altered in ZO-1-depleted cells (15, 16). NMIIC was found to form a sarcomeric belt that colocalized with tight junctions in epithelial cells in some organs, suggesting the interaction between NMIIC and tight junction even *in vivo* (17). Finally, NMIIA was reported to destroy the tight junction in oxygen and glucose-deprived mouse brain endothelial cells (18). Therefore, we hypothesized that if NMIIs are expressed in the lacrimal gland, they may play a role in the regulation of tight junction proteins, and thus modulate tear secretion, providing evidence for a paracellular water pathway. To test this hypothesis, we first determined that NMIIC was expressed at apical junctions in ductal cells of NMIIC-GFP knockin mice. We then used both genetic and pharmacological perturbations of NMIIC to investigate a putative role regulating the tear secretion. In addition, we found, intriguingly, that the lacrimal gland and the salivary gland employ distinct mechanisms of water secretion, with negligible paracellular water permeability between ductal cells in the latter, even though the molecular make-up of acinar cells and ductal cells is similar in both organs. Our data provides new mechanistic insights into paracellular water secretion *via* ductal cell tight junctions in the murine lacrimal gland but not in the parotid gland.

## Results

### 1. NMIIC localizes to the apical junctions of ductal cells in murine lacrimal gland

Myosin is a hexamer consisting of 2 heavy chains and 4 light chains. The three different NMII paralogs, NMIIA, NMIIB or NMIIC, differ in their heavy chains. To determine which are expressed in the murine lacrimal gland, we investigated their expression levels in our previous RNA-seq data (accession number DRA010121 at DDBJ Sequenced Read Archive) (19). We determined that the predominant heavy chains in the lacrimal gland are Myh9 (NMIIA), and Myh14 (NMIIC), while Myh10 (NMIIB) expression level was substantially lower (Figure 1A; 8.59 ± 2.18 in NMIIA, 0.12 ± 0.03 in NMIIB, and 2.12 ± 0.30 in NMIIC). Myh11 was also expressed as a smooth muscle myosin-heavy chain (Figure 1A; 3.11 ± 1.02 in Myh11), suggesting that it is expressed in myoepithelial cells in the lacrimal gland. Carbachol, a muscarinic receptor agonist, is well-established to induce regulated exocytosis in lacrimal acinar cells, facilitated by actin and NMII (20). Consistent with this, immunofluorescence staining of NMIIA validated that NMIIA was expressed in acinar cells, where it localized to secretory granules fused to the apical membrane during carbachol-induced exocytosis (Figure 1B). This NMIIA coat on exocytic granules in the lacrimal gland is similar to what was reported in the salivary gland (21), suggesting the NMIIA has a role in extruding granule contents across exocrine glands.

**Figure 1.**
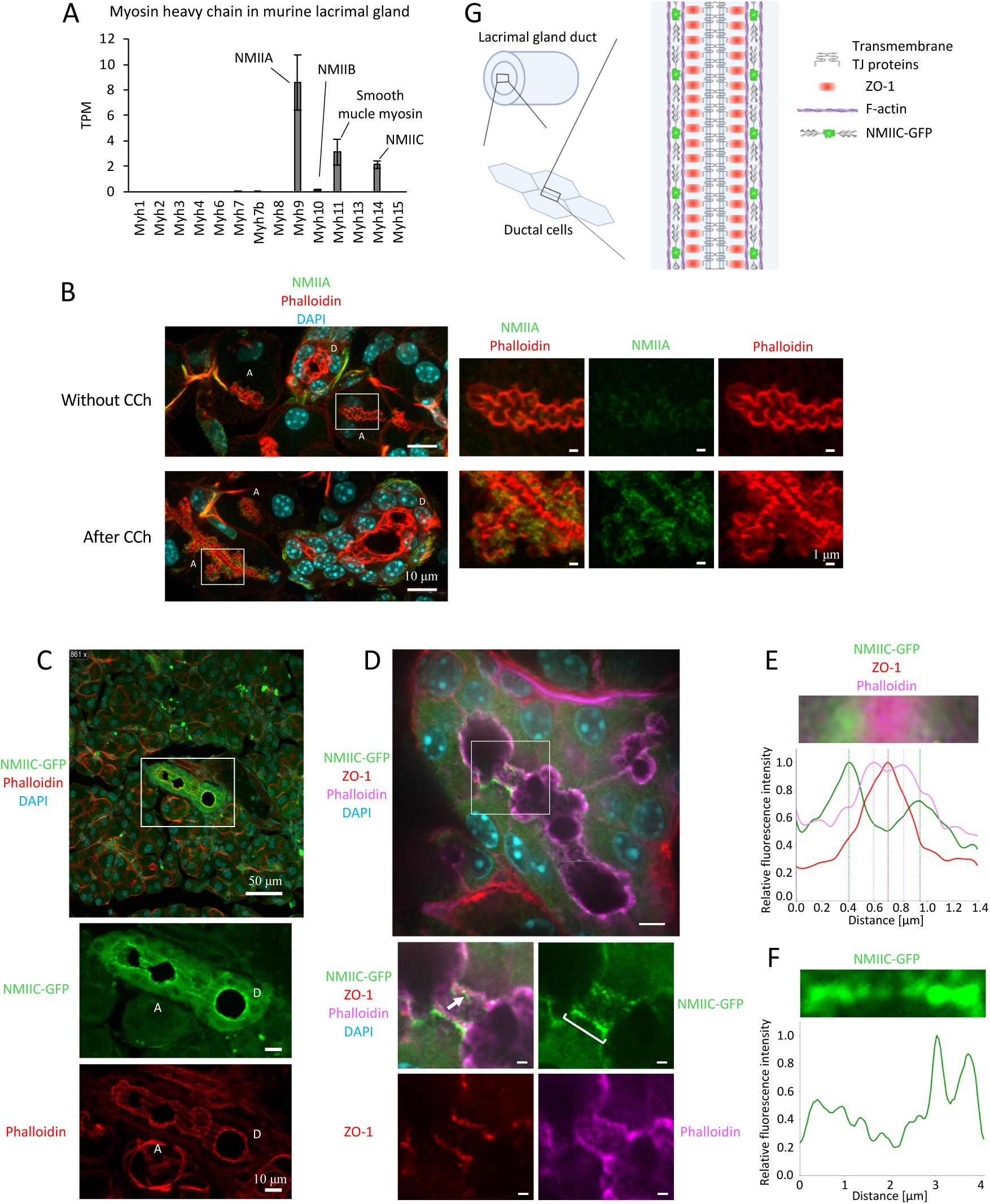
NMIIC is enriched at tight junctions between ductal cells in murine lacrimal gland. (A) Myosin heavy chain (Myh) mRNA expression level in the murine lacrimal gland from previous RNA-seq data (accession number DRA010121 at DDBJ Sequenced Read Archive) (19). (B) Localization of NMIIA in the lacrimal gland from wildtype mice without or after carbachol (CCh) stimulation. NMIIA (green) localizes on the membrane of secretory granules in acinar cells after CCh stimulation. Phalloidin (red) labels F-actin. DAPI (blue) labels the nucleus. Scale bars, 10μm in the left images and 1 μm in the right images. (C) Localization of NMIIC in the lacrimal gland of NMIIC-GFP mice. NMIIC-GFP (green) localizes at the apical junctions of ductal cells in the lacrimal gland. Phalloidin (red) labels F-actin. DAPI (blue) labels the nucleus. D; duct, A; acinar cells. Scale bars, 50 μm in the upper image and 10 μm in the lower images. (D) Localization of NMIIC at high magnification. NMIIC-GFP (green) was expressed as puncta along junstional F-actin (magenta) and ZO-1 (red). DAPI (blue) represents the nucleus. Scale bars, 5 μm in the upper image and 1 μm in the lower images. (E) Fluorescence intensity of NMIIC-GFP, ZO-1 and Phalloidin at the arrow in Figure 1D, crossing the junction. (F) Fluorescence intensity of NMIIC-GFP along the bracket in Figure 1D. (G) Schematic of the ductal junction of the lacrimal gland. Created with BioRender.com.

We next determined the localization of NMIIC in lacrimal glands from NMIIC-GFP mice. Distinct from the localization of NMIIA, NMIIC-GFP was enriched at the apical junction of ductal cells and was expressed in acinar cells at much lower levels (Figure 1C). When observed at higher magnification, we determined that NMIIC at the ductal cell apical junctions was expressed as regularly-spaced puncta along F-actin, (Figure 1D and 1F). Detailed analysis of fluorescence intensity (FI) across the apical junction of ductal cells indicated that FI peaks of F-actin and NMIIC flanked and overlapped partially, with that of ZO-1 (Figure 1E), suggestive of some level of interaction between these proteins (Figure 1G).

### 2. Carbachol-induced tear volume is increased in NMIIC-KO mice via tight junction disruption

To determine whether NMIIC is involved in tear secretion, we measured tear volume in NMIIC-KO mice and wild-type mice by stimulating surgically exposed lacrimal glands with carbachol. Tear volume before carbachol stimulation, i.e., basal tear volume, did not show a significant difference between these groups (Figure 2A-ii; wild-type vs. NMIIC-KO = 3.1 ± 1.1 vs. 2.5 ± 0.9 mm/3 min). However, NMIIC-KO mice showed significantly increased tear secretion after carbachol stimulation compared to wild-type mice (Figure 2A-ii; wild-type vs. NMIIC-KO = 15.5 ± 5.8 vs. 21.0 ± 3.8 mm/3 min). The incremental tear volume, which was calculated by subtracting tear volume before carbachol stimulation from tear volume after carbachol stimulation, also showed a significant increase in the NMIIC-KO group (Figure 2A-iii; wild-type vs. NMIIC-KO = 12.4 ± 5.8 vs. 18.5 ± 3.3 mm/3 min).

**Figure 2.**
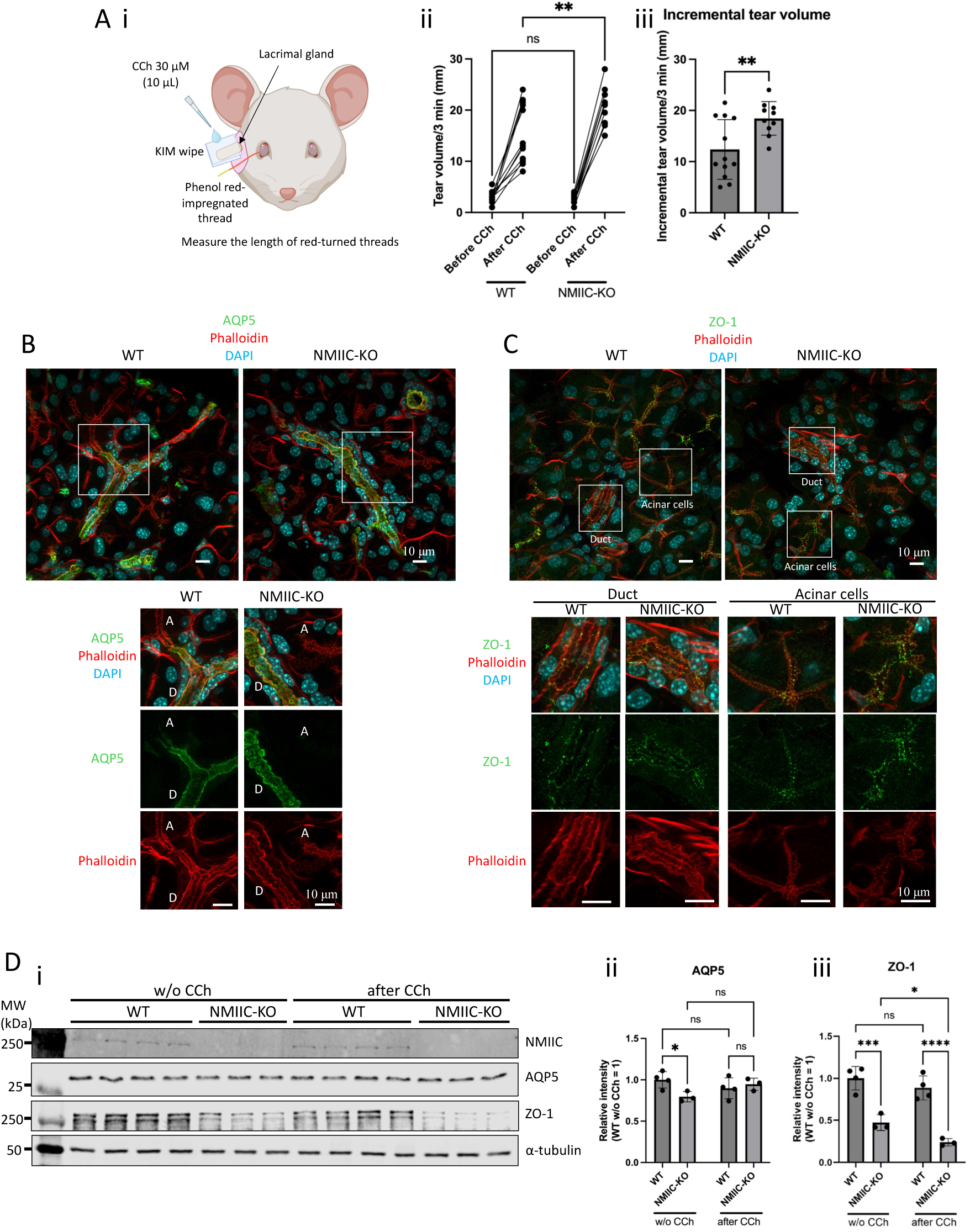
CCh-induced tear volume increased and ZO-1 expression level reduced in NMIIC KO mice. (A-i) Illustration of procedure for tear volume measurement. The lacrimal gland of anesthetized mouse was surgically exposed wrapped with a KIM wipe upon which CCh was applied. A phenol red-impregnated thread was placed at the canthus of eye for 3 min each before and after CCh stimulation, and tear volume was measured as the length of the thread that turned red due to exposure to moisture. Created with BioRender.com. (A-ii) Tear volume from NMIIC-KO mice and WT mice. (A-iii) Incremental tear volume from NMIIC-KO mice and WT mice, calculated by subtracting tear volume before CCh stimulation from tear volume after CCh stimulation. \*\**P* < 0.01. Two-way ANOVA followed by Sidak’s multiple comparison test (A-ii) or *t*-test (A-iii). N = 10-12. (B) Localization of AQP5 (green) in lacrimal gland from NMIIC-KO mice and WT mice. Phalloidin (red) represents F-actin. DAPI (blue) represents the nucleus. D; duct, A; acinar cells. Scare bars, 10 μm. (C) Localization of ZO-1 (green) in lacrimal gland from NMIIC-KO mice and WT mice. Phalloidin (red) represents F-actin. DAPI (blue) represents the nucleus. Scare bars, 10 μm. (D-i) Western blot of NMIIC, AQP5, ZO-1, and α-tubulin in lacrimal gland from NMIIC-KO mice and WT mice, after CCh stimulation to the lacrimal gland or without CCh. (D-ii) Quantification of AQP5 band intensity. (D-iii) Quantification of ZO-1 band intensity. \**P* < 0.05, \*\**P* < 0.01, \*\*\**P* < 0.001, \*\*\*\**P* < 0.0001. Two-way ANOVA followed by Fisher’s LSD test. N = 3 or 4.

We also investigated the localization of AQP5, a dominant water channel in the lacrimal gland, in NMIIC-KO lacrimal glands by immunofluorescence staining. AQP5 was mainly expressed on the apical membrane of the duct, and present at lower levels on the apical membrane of acinar cells in lacrimal gland of wild-type mice. This expression pattern was not affected in NMIIC-KO mice, suggesting that NMIIC does not affect the localization of AQP5 (Figure 2B). Next, we investigated the localization of ZO-1, one of the major tight junction scaffold proteins. ZO-1 was expressed at the apical junctions of both ductal cells and acinar cells. NMIIC loss did not affect the localization of ZO-1 (Figure 2C).

Finally, we determined the protein levels of AQP5 and ZO-1 in lacrimal glands from NMIIC-KO and wildtype without and after carbachol stimulation. NMIIC knockout was also confirmed since there was no band in the NMIIC-KO group (Figure 2D-i). AQP5 protein level showed only a slight decrease in the NMIIC-KO lacrimal gland without carbachol stimulation and remained unchanged after carbachol stimulation (Figure 2D-i and 2D-ii; wild-type vs. NMIIC-KO = 1.00 ± 0.10 vs. 0.80 ± 0.06 without carbachol stimulation, and 0.90 ± 0.13 vs. 0.95 ± 0.07 after carbachol stimulation). Conversely, ZO-1 was notably reduced by 53% and 73% in the NMIIC-KO lacrimal gland compared to wild-type, without and after carbachol stimulation, respectively (Figure 2D-i and 2D-iii; wild-type vs. NMIIC-KO = 1.00 ± 0.14 vs. 0.47 ± 0.10 without carbachol stimulation, and 0.89 ± 0.14 vs. 0.24 ± 0.04 after carbachol stimulation). These data suggested that the mechanism underlying the increase in carbachol-induced tear volume in mice lacking NMIIC involves reduced ZO-1 expression.

### 3. Tears from NMIIC-KO mice showed brighter fluorescence when a fluorescent dye was mixed with carbachol

Given these data, we speculated that leakage between ductal cells might lead to increased tear secretion caused by the loosening of the tight junction. To confirm this possibility, we performed a tear secretion experiment again, but this time, the carbachol solution used to stimulate tear secretion also contained fluorescein—a fluorescence compound with a small molecular weight, which is hydrophilic and thus impermeable to cell membrane. We then collected tears from NMIIC-KO and wild-type mice simultaneously on a nitrocellulose membrane and observed them under the microscope (Figure 3A-i). Tears from NMIIC-KO mice appeared more strongly fluorescent compared to wild-type mice (Figure 3A-ii). The quantification of fluorescence intensity from NMIIC-KO tears confirmed a significant increase compared to wild-type mice (Figure 3A-iii; wild-type vs. NMIIC-KO = 1.00 ± 0.00 vs. 3.26 ± 1.41). These data provide evidence for leakage between cells in the NMIIC-KO lacrimal gland.

**Figure 3.**
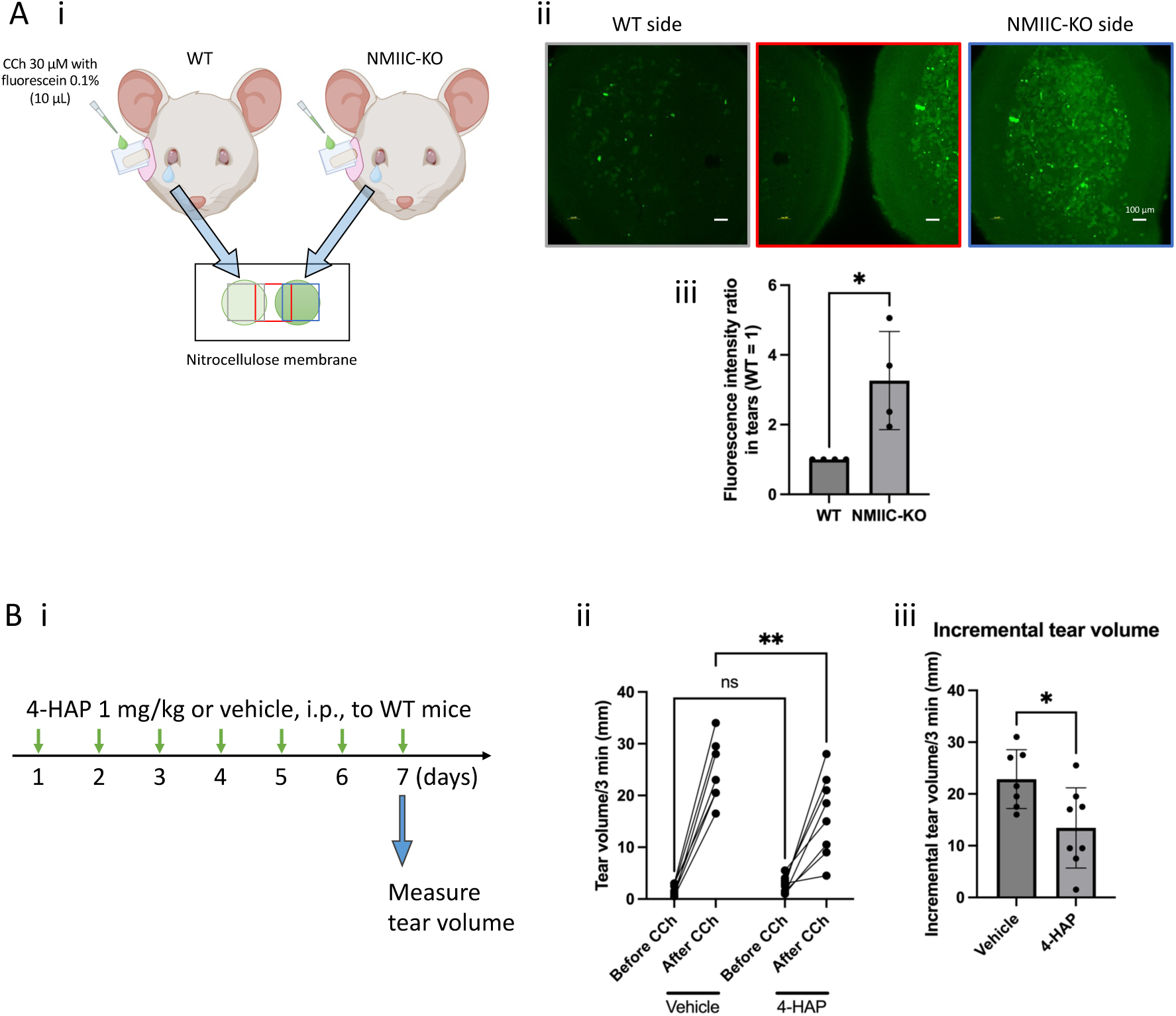
Evidence for paracellular water secretion in the lacrimal glands. (A) NMIIC-KO mice showed higher fluorescence in secreted tears when the lacrimal gland was stimulated with CCh mixed with fluorescein. (A-i) Illustration of collection of tears after lacrimal gland stimulation with CCh-fluorescein mixture. Tears were collected and put on the nitrocellulose membrane and then observed under the microscope. Created with BioRender.com. (A-ii) Images of tear collected on nitrocellulose membrane. Scale bars, 100 μm. (A-iii) Quantification of relative fluorescence intensity in the same field of view of microscope. \**P* < 0.05. Unpaired *t*-test with Welch’s correction. N = 4. (B) Impact of NMIIC activator, 4-HAP, on tear volume induced by CCh. (B-i) Protocol of 4-HAP treatment. 4-HAP was administered at 1 mg/kg, i.p., to WT mice once a day for 7 days before measuring tear volume. (B-ii) Tear volume from 4-HAP-treated mice and vehicle-treated mice. (B-iii) Incremental tear volume from 4-HAP-treated mice and vehicle-treated mice, calculated by subtracting tear volume before CCh stimulation from tear volume after CCh stimulation. \**P* < 0.05, \*\**P* < 0.01. Two-way ANOVA followed by Sidak’s multiple comparison test (B-ii) or *t*-test (B-iii). N = 7 or 8.

### 4. Activation of NMIIC by 4-HAP reduces carbachol-induced tear volume

For further confirmation of a role for NMIIC in regulating tear secretion, we administered 4’-hydroxyacetophenone (4-HAP), an NMIIC-activator, to WT mice for 7 days based on a previous study (22), and then measured tear volume (Figure 3B-i). 4-HAP-injected mice showed reduced tear volume after carbachol stimulation compared to vehicle-injected mice (Figure 3B-ii; wild-type vs. NMIIC-KO = 24.6 ± 6.1 vs. 16.2 ± 7.9 mm/3 min), while there was no significant difference between these two groups before carbachol stimulation (Figure 3B-ii; wild-type vs. NMIIC-KO = 1.7 ± 1.1 vs. 2.8 ± 1.6 mm/3 min). Incremental tear volume in 4-HAP-injected mice also showed a significant increase compared to vehicle-injected mice (Figure 3B-iii; wild-type vs. NMIIC-KO = 22.9 ± 5.7 vs. 13.4 ± 7.7 mm/3 min).

We then injected NMIIC-GFP mice with 4-HAP or vehicle for 7 days and performed immunofluorescence staining of AQP5 and ZO-1 to determine whether the localizations of these proteins or NMIIC are affected. NMIIC, AQP5 and ZO-1 localization did not change between the 4-HAP and vehicle groups (Supplemental Figure S1A and 1B). We also investigated the protein expression levels of AQP5 and ZO-1 by western blot. The results showed that the NMIIC-activation by 4-HAP did not change AQP5 and ZO-1 expression levels (Supplemental Figure S1C). In summary, pharmacological activation of NMIIC reduced tear secretion but did not affect localization or expression level of AQP5 or ZO-1.

### 5. NMIIC loss reduces water secretion in salivary gland

To gain further insight on NMIIC-mediated regulation of tight junctions in exocrine glands we investigated the role of NMIIC in saliva secretion in the parotid gland and compared it with the lacrimal gland data. The expression level of NMIIC in parotid gland is higher than that in lacrimal gland (Figure 4A-i and 4A-ii; lacrimal gland vs. parotid gland =1.00 ± 0.20 vs. 4.09 ± 1.07). NMIIC-GFP fluorescence was enriched at the apical junction of ductal cells and was less expressed in acinar cells in the parotid gland (Figure 4B-i). The localization and pattern of NMIIC, in puncta, was similar to that of NMIIC in the lacrimal gland (Figure 4B-ii). As we did in the lacrimal gland, we measured saliva volume in NMIIC-KO mice and wild-type mice by stimulating the exposed parotid gland with carbachol (Figure 4C-i). Strikingly, contrary to the tear secretion data, NMIIC-KO mice showed decreased saliva secretion after carbachol stimulation compared to wild-type mice (Figure 4C-ii and 4C-iii; wild-type vs. NMIIC-KO = 60.2 ± 11.3 vs. 40.2 ± 3.5 mg/15 min).

**Figure 4.**
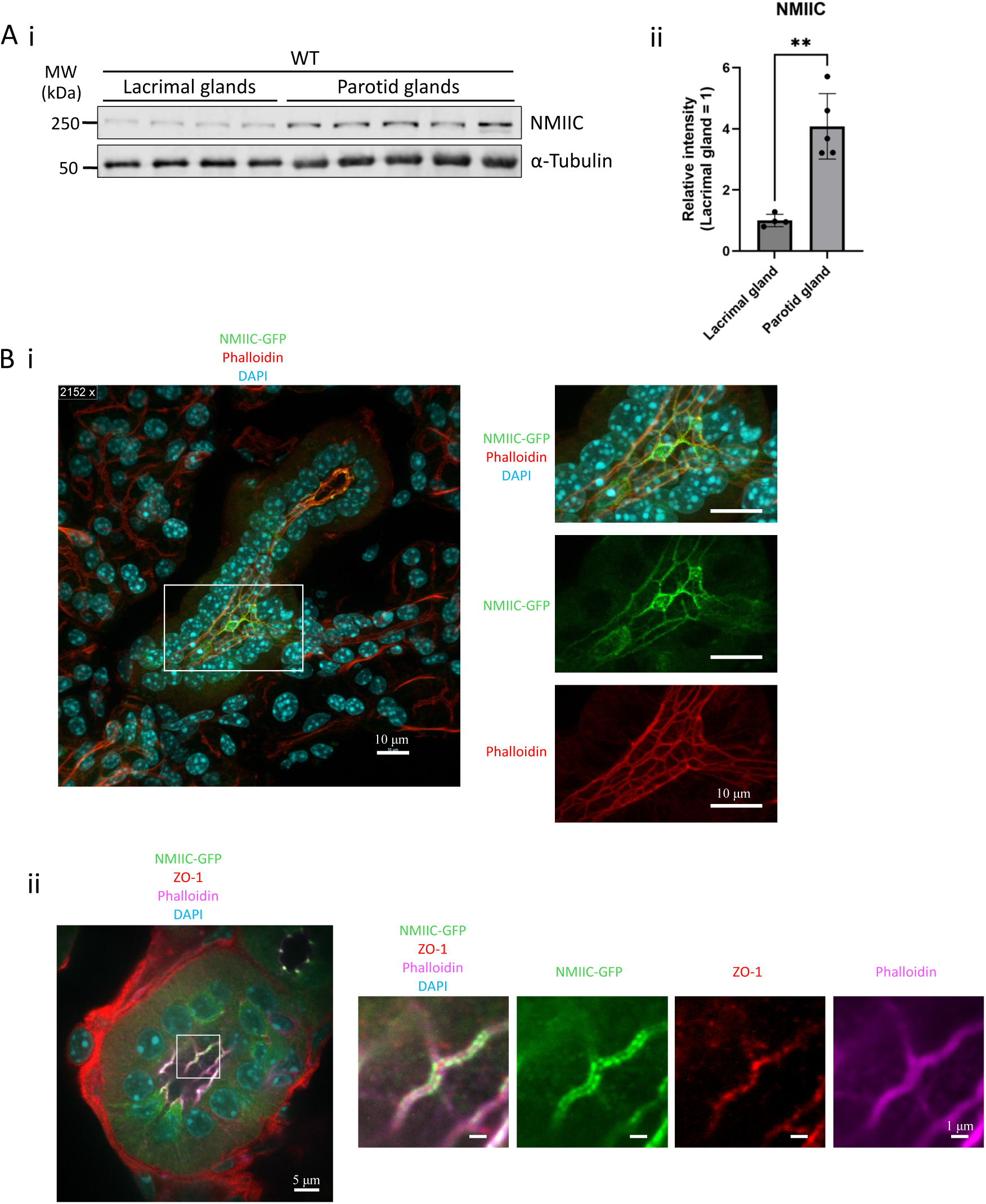
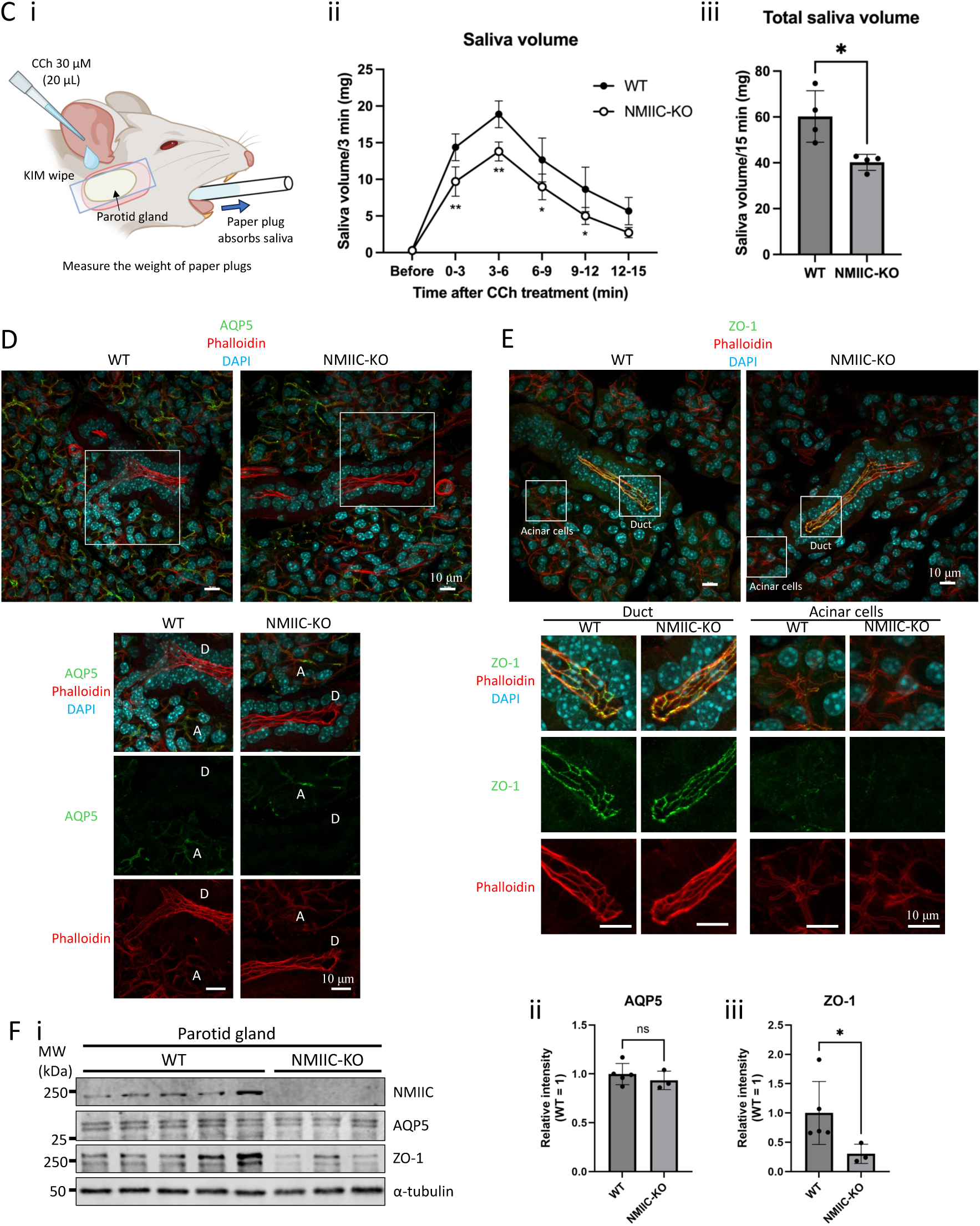
CCh-induced saliva volume and ZO-1 expression level are reduced in NMIIC KO mice. (A-i) Comparison of NMIIC expression level between lacrimal glands and parotid glands by western blot. (A-ii) Quantification of NMIIC band intensity. \*\**P* < 0.01. Student’s *t*-test. N = 4 or 5. (B-i) Localization of NMIIC in parotid gland of NMIIC-GFP. NMIIC-GFP (green) localizes at the apical junction in ductal cells in the parotid gland from NMIIC-GFP mice. Phalloidin (red) represents F-actin. DAPI (blue) represents the nucleus. Scale bars, 10 μm. (B-ii) Localization of NMIIC at higher magnification. NMIIC-GFP (green) was expressed as puncta along junctional F-actin (magenta) and ZO-1 (red). DAPI (blue) represents the nucleus. Scale bars, 5 μm in the left image and 1 μm in the right images. (C) Saliva secretion in NMIIC-KO mice. (C-i) Illustration of procedure for saliva volume measurement. The parotid gland of anesthetized mouse was surgically exposed wrapped with a KIM wipe upon which CCh was applied. Rolled paper was inserted into oral cavity before and after CCh stimulation every 3 min up to 15 min, and saliva volume was measured as the weight of absorbed saliva. Created with BioRender.com. (C-ii) Time-dependent change of saliva volume from NMIIC-KO mice and WT mice. (C-iii) Total saliva volume for 15 min after CCh stimulation from NMIIC-KO mice and WT mice. \**P* < 0.05, \*\**P* < 0.01. Two-way ANOVA followed by Sidak’s multiple comparison test (C-ii) or *t*-test (C-iii). N = 4. (D) Localization of AQP5 (green) in parotid gland from NMIIC-KO mice and WT mice. Phalloidin (red) represents F-actin. DAPI (blue) represents the nucleus. D; duct, A; acinar cells. Scare bars, 10 μm. (E) Localization of ZO-1 (green) in parotid gland from NMIIC-KO mice and WT mice. Phalloidin (red) represents F-actin. DAPI (blue) represents the nucleus. Scare bars, 10 μm. (F-i) Western blot of NMIIC, AQP5, ZO-1, and α-tubulin in parotid gland from NMIIC-KO mice and WT mice without CCh stimulation. (F-ii) Quantification of AQP5 band intensity. (F-iii) Quantification of ZO-1 band intensity. \**P* < 0.05. Student’s *t*-test. N = 3-5.

Distinct from the lacrimal gland, immunofluorescence staining showed that AQP5 was mainly expressed on the apical membrane of the acinar cells, and not expressed at apical junctions of ductal cells in wild-type lacrimal gland. This localization was not affected in the NMIIC-KO mice (Figure 4D). ZO-1 was enriched at the apical junctions between ductal cells, and at a lower level between acinar cells. NMIIC-KO did not affect the localization of ZO-1 (Figure 4E).

Finally, we performed western blots to check the protein level of AQP5 and ZO-1 in the parotid gland from NMIIC-KO (Figure 4F-i) and wildtype. Two AQP5 bands were detected in parotid glands, differing from a single band in the lacrimal glands. The two bands of AQP5 in the parotid gland are likely glycosylated and non-glycosylated forms previously reported in salivary glands (23). Similar to lacrimal gland data, ZO-1 was reduced by 70% in the NMIIC-KO parotid gland compared to wild-type (Figure 4F-i and 4F-iii; wild-type vs. NMIIC-KO = 1.00 ± 0.54 vs. 0.30 ± 0.16), whereas AQP5 remained unchanged (Figure 4F-i and 4F-ii; wild-type vs. NMIIC-KO = 1.00 ± 0.11 vs. 0.93 ± 0.10). While NMIIC loss resulted in a significant reduction in ZO-1 expression in both lacrimal and parotid glands, the opposing consequences of increased tear volume but reduced saliva volume likely reflect distinct mechanisms of secretion and degrees of paracellular water permeability between these organs.

## Discussion

In this study, we investigated the presence of a paracellular water secretion pathway in tear secretion by targeting NMIIC, which was expressed at apical junctions of ductal cells in the murine lacrimal gland, overlapping with tight junctions. We found that tear volume after carbachol stimulation was significantly increased in NMIIC-KO mice. Moreover, when we mixed a fluorescent dye with carbachol, tears from NMIIC-KO mice showed brighter fluorescence. Administration of an NMIIC activator, 4-HAP, to wildtype mice, inhibited tear secretion. The localization of AQP5 and the tight junction protein ZO-1 in lacrimal glands by immunofluorescence were unchanged in NMIIC-KO mice. However, ZO-1 protein levels were significantly reduced in NMIIC-KO lacrimal glands, while AQP5 levels remained unchanged. Intriguingly, contrary to tear secretion, carbachol-induced saliva secretion was reduced in NMIIC-KO mice, even with reduced ZO-1 expression. In summary, our findings reveal a new mechanism for paracellular water secretion in the lacrimal gland *via* NMIIC-mediated regulation of ductal cell tight junctions, which seems not to exist in the parotid gland (as illustrated schematically in Figure 5).

**Figure 5.**
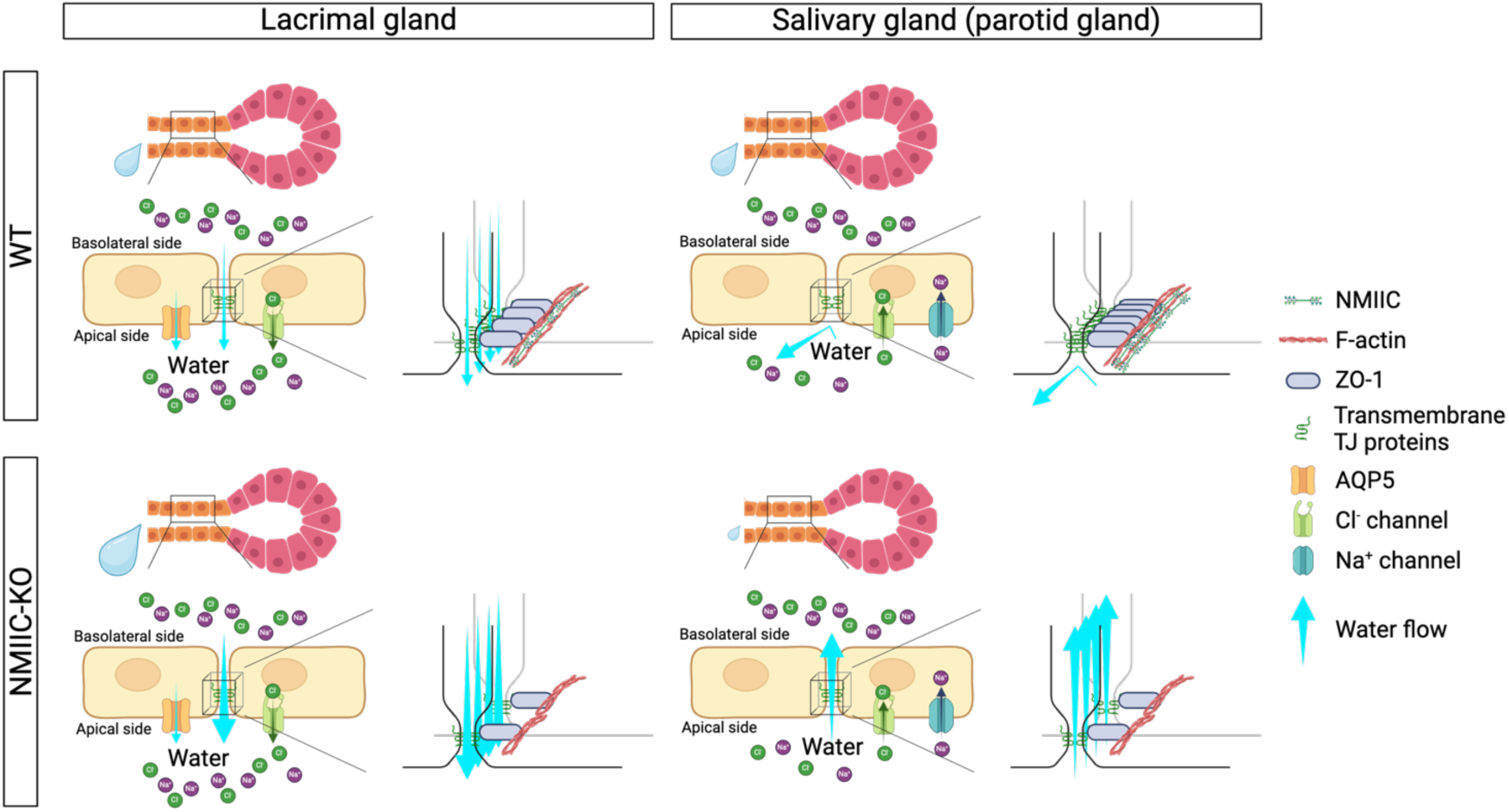
Model for NMIIC-mediated regulation of paracellular water transport in tear and saliva secretion in mice. NMIIC is enriched at ductal cell tight junctions in the murine lacrimal and salivary glands (parotid glands). Loss of NMIIC results in reduced expression of ZO-1 in both glands. In the lacrimal gland, water flows through AQP5 channels and tight junctions towards the lumen due to higher ion concentration and positive osmotic pressure. Ablation of NMIIC, and consequential reduction of ZO-1 and loosening of tight junctions causes increased water flow and thus tear secretion increases. Conversely, in the salivary gland ductal cells do not express water channels such as AQP5. Thus, while ions are pumped back into ductal cells from the lumen, water remains, resulting in a negative osmotic pressure in the lumen. In this case when tight junctional permeability is increased in absence of NMIIC, water flows out of the lumen resulting in reduced saliva secretion. Created with BioRender.com.

### 1. Water secretion in duct in the lacrimal gland

The ducts in the lacrimal gland are reported to secrete water (24). In the murine lacrimal gland, the dominant water channel for transcellular water secretion is thought to be AQP5 based on previous reports (25) as well as to our previous RNA-seq data (accession number DRA010121 at DDBJ Sequenced Read Archive) (19). Recently, single-cell RNA-seq data revealed that AQP5 is more highly expressed in ductal cells compared to acinar cells in the murine lacrimal gland (26). This is consistent with our data showing that ductal cells have higher AQP5 expression in their apical membrane than acinar cells (Figure 2B), and also consistent with previous papers in which AQP5 was strongly stained in ducts compared to acini (27, 28). The single-cell RNA-seq data also suggested that ductal cells have several ion channels and ion transporters as well as Na+-K+ ATPase, which have lower, or no expression in acinar cells (26). This also suggests that the lacrimal ducts are responsible for water secretion since the secreted ions create higher osmotic pressure in the ductal lumen side than in extracellular fluid or cytosol in ductal cells and make water move into the lumen side of the duct. Therefore, the possibility of paracellular water secretion in the “duct” is also reasonable. Acinar cells in the lacrimal gland seem to be responsible for protein secretion *via* exocytosis when the muscarinic receptor is stimulated by carbachol (Figure 1B) (29–31), although we do not know whether the water secretion also occurs in acinar cells.

### 2. Possibility of paracellular water secretion from between ductal cells in the lacrimal gland

NMIIC-KO mice showed increased tear secretion (Figure 2A) and 4-HAP-injected mice showed decreased tear secretion (Figure 3B). Moreover, after stimulating the lacrimal gland with a fluorescein-containing carbachol solution, tears from NMIIC-KO mice were significantly more fluorescent than wild-type (Figure 3A). Since fluorescein, a fluorescence compound with a small molecular weight, is hydrophilic and thus impermeable to cell membranes, these data suggest either: 1) paracellular water secretion is upregulated/downregulated depending on NMIIC status, or 2) the endocytosis of extracellular fluid from basolateral side and exocytosis of the granules into the lumen side are upregulated/downregulated depending on NMIIC status. However, since ZO-1 expression levels are reduced in NMIIC-KO mice (Figure 2D) and the expression levels of NMIIC in the acinar cells, where much higher exocytosis activity exists (Figure 1B and 1C) (32), is significantly lower than in ductal cells, the first hypothesis is more plausible.

### 3. Different paracellular water permeability of duct in lacrimal gland and salivary gland

There is currently no study that reports evidence in support of the existence of the paracellular water pathway in tear secretion. On the other hand, there are some reports regarding this pathway in the salivary gland. Murakami et al. reported the possibility of paracellular water transport based on a mathematical model fitting to the experimental permeability data (33) and exposure to a hypertonic solution (34) in perfused rat submandibular glands. Experiments analyzing the perfusion of lucifer yellow, a cellular impermeable fluorescence substance, with some saliva stimulants, also support the presence of a paracellular water pathway in saliva secretion (35–37). Therefore, it is not unlikely that paracellular water secretion occurs in the lacrimal gland. However, a major distinction highlighted by our study is that paracellular water secretion in the salivary gland is considered to occur between acinar cells (37–39), because saliva secreted into the oral cavity is known to be hypotonic after reabsorption of ions from primary saliva in the duct (40), and thus the junction of duct should be sealed tightly. If paracellular water permeability were high in the duct in the salivary gland, then saliva would become close to isotonic fluid during the flow of primary fluid from acini towards the duct. We hypothesize this to be the mechanism underlying reduced saliva secretion in absence of NMIIC (Figure 4C), specifically that NMIIC-KO loosened the tight junction *via* reduced ZO-1 expression level (Figure 4F) causing water to be reabsorbed through the ductal junctions due to osmotic pressure differences between hypotonic primary saliva and isotonic extracellular fluid. Our data thus revealed that paracellular water secretion occurs in acinar cells but not in ducts in the healthy salivary gland.

On the other hand, tears are reported to be isotonic fluid (41), supporting the likelihood of paracellular water transport in the duct. These differences are also supported by the opposite localization of ion/water channels such as chloride channels and AQP5 as well as ion transporters such as NKCC1; they are enriched in ductal cells in the lacrimal gland (Figure 2B) (26, 42) but in acinar cells in the salivary gland (Figure 4D) (43). Therefore, the paracellular water permeability in ducts is considered to be high in the lacrimal gland but quite low in the salivary gland.

### 4. Mechanism of NMIIC-mediated regulation of paracellular water secretion in the lacrimal gland

4-HAP is known to activate NMIIB and NMIIC (44), but it is considered to be exclusively effective on NMIIC in this study because NMIIC mRNA is expressed in the murine lacrimal gland but NMIIB mRNA expression levels are negligible (Figure 1A).

Activation by 4-HAP causes an increase in NMIIC assembly in cells. Based on this fact, we propose two possibilities: one is that 4-HAP-treatments results in increased NMIIC-forces on the actomyosin belt that overlaps with the tight junction, thus increasing barrier integrity and reducing tear secretion. The other possibility is that activating NMIIC enables the recruitment of tight junction proteins, thus influencing tear volume depending on the tight junction expression level. Our results revealed that the former is more likely as there was no significant difference in ZO-1 expression levels after 4-HAP treatment (Supplemental Figure S1C). However, we consider the latter also could be possible if 4-HAP was administered to mice for a longer duration because several studies report that changing forces on tight junctions can alter their composition and function; e.g., physiological shear stress was reported to increase the expression level of tight junctions in brain microvascular endothelial cells (45), and junctional tension was also reported to lead to the transportation of non-junctional ZO-1 toward tight junctions in zebrafish embryos (12). Furthermore, myosin light chain is reported to be related to forming tight junctions (46). These reports provide mechanistic insight into NMIIC-regulating tightening of tight junctions which leads to changing volume of paracellular water transport, although further study is needed to confirm this.

### 5. Translational perspective of this study

Dry eye is a condition that continues to increase in prevalence, especially as tear volume in humans tends to decrease with aging (2, 47). In reporting NMIIC as a direct modulator of water secretion through a paracellular pathway in ductal cells of the lacrimal and salivary glands, we have identified a promising target for the development of pharmaceutical therapeutics for dry eye or dry mouth.

### 6. Conclusion

Our findings are the first to provide evidence for the involvement of a paracellular water pathway in tear secretion. Furthermore, we find that this pathway is regulated by NMIIC *via* modulation of tight junctions in the ductal cells in the lacrimal glands and salivary glands. This study also highlights NMIIC as a potential target for the development treatments for dry eye/mouth syndrome.

## Materials and Methods

### 1. Materials

Phenol red-impregnated thread (Zone-Quick^®^) was purchased from Ayumi Pharmaceutical Co. (Tokyo, Japan). Medetomidine (Placadine^®^) was purchased from Modern Vet (Springfield, TN, USA), midazolam was from Hospira (Lake Forest, IL, USA), butorphanol (Torphadine^®^) was from Dechra Pharmaceuticals plc (Northwich, UK), paper plugs (JM paper point^®^) were from J. Morita Corp. (Osaka, Japan), RIPA lysis buffer, protease inhibitor (Pierce Protease Inhibitor Tablets, EDTA-free), and optical cutting temperature (O.C.T.) coumpound were from Thermo Fisher (Waltham, MA, USA), nitrocellulose membrane and Triton X-100 were from Bio-Rad Laboratories Inc (Hercules, CA, USA), skim milk was from BD Life Sciences (Franklin Lakes, NJ, USA), 4’-hydroxyacetophenone (4-HAP) was from Sigma-Aldrich (St. Louis, MO, USA), 10% normal goat serum, rabbit anti-NMIIC antibody, goat anti-rabbit IgG antibody conjugated with Alexa Fluor^®^ 488 or Alexa Fluor^®^ 568, goat anti-rat IgG antibody conjugated with Alexa Fluor^®^ 488 or Alexa Fluor^®^ 568, phalloidin conjugated with Alexa FluoroTM 568 or Alexa FluoroTM 647, and mounting media (Fluoromount-GTM with DAPI) were from Invitrogen (Carlsbad, CA, USA), rabbit anti-NMIIA antibody was from BioLegend (San Diego, CA, USA), rabbit anti-AQP5 antibody was from Alomone Labs (Jerusalem, Israel), rat anti-ZO-1 antibody was from BiCell Scientific (Maryland Heights, MO, USA), rabbit anti-ZO-1 antibody was from GeneTex (Irvine, CA, USA), mouse anti-α-tubulin antibody was from Santa Cruz Biotechnology (Dallas, TX, USA), anti-mouse IgG antibody conjugated with Alexa Fluor^®^ 680 and anti-rabbit IgG antibody conjugated with Alexa Fluor^®^ 790 were from Jackson ImmunoResearch Inc. (West Grove, PA, USA).

### 2. Mice

NMIIC-KO and NMIIC-GFP mice were generated as described in (48) and (17), respectively. C57BL/6J mice were used as a control for the NMIIC-KO mice. All mice used in the experiments were male and 2 months old (8-13 weeks old).

All mice were maintained under controlled conditions (20 ± 2°C, 50 ± 20% humidity, 12 h light/dark cycle) in the animal facility at the University of Virginia. All mice were given free access to water and standard chow (Teklad LM-485 mouse/rat sterilizable diet; Inotiv; West Lafayette, IN, USA).

All animal experiments were performed according to protocols approved by the animal care and use committee (ACUC) at the University of Virginia (Approval Numbers: 4382). All efforts were made to minimize suffering of the animals.

### 3. Myh expression analysis from mRNA-seq data

mRNA-seq analysis data that we performed before (19) was utilized for the mRNA expression analysis for the myosin heavy chain (Myh) in the murine lacrimal gland. The expression profile was calculated as transcripts per million (TPM). The nucleotide sequence data are available in the DDBJ Sequenced Read Archive under accession number DRA010121.

### 4. Measurement of tear volume

The tear volume was determined by the cotton thread test using a phenol red-impregnated thread. Mice were anesthetized by intraperitoneal (i.p.) injection of an anesthetic agent mixture (0.75 mg/kg of medetomidine, 4.0 mg/kg of midazolam and 5.0 mg/kg of butorphanol) at a volume of 0.05 mL/10 g body weight. The anesthetic agent mixture was decided based on previous reports (19, 49, 50). The mice were placed on a heating pad and lacrimal glands were exposed under the anesthesia, and then wrapped with a small cut of cellulose mesh (Kimtech; Kimberly-Clark, Irving, TX, USA). Carbachol solution was applied topically on the lacrimal glands at the concentration of 0.3 μM and at a volume of 10 μL, which was decided based on our results of the tear volume measurement induced by the different concentration of carbachol as well as a previous paper (51). The volume of tear fluid was measured by carefully placing a phenol red-impregnated thread at the canthus of each eye for 3 min before and after carbachol stimulation. The length of thread that changed color due to absorption of tear fluid was measured in millimeters. The incremental tear volume was calculated by subtracting the tear volume before carbachol stimulation from the tear volume after carbachol stimulation.

### 5. Measurement of saliva volume

The saliva volume was determined by a gravimetric method using paper. With the animal under anesthesia as described above, mice were placed on a heating pad and parotid glands were exposed, and then a small cut of cellulose mesh were placed on the parotid glands. Carbachol solution was applied topically on the parotid glands at the concentration of 0.3 μM and at a volume of 20 μL. Then the secreted saliva was absorbed into paper plugs inserted into the oral cavity and exchanged at 3 min intervals up to 15 min after starting carbachol stimulation. The saliva-saturated plugs were weighed and corrected for the original weight of the paper plug. The volume of secreted saliva was calculated as the increase in weight of each paper plug. The total saliva volume was calculated by summing up the increase in weight of each paper plug after carbachol stimulation.

### 6. Immunofluorescence staining

The lacrimal and parotid glands were isolated under the anesthesia as described above. Lacrimal glands and parotid glands were isolated, and then the glands were fixed in 4% paraformaldehyde in phosphate buffered saline (PBS; pH 7.4) for 1 h at room temperature for the lacrimal glands or overnight at 4℃ for the parotid glands. After fixation, the lacrimal glands and parotid glands were put through a sucrose gradient (10% → 20% → 30%) for cryoprotection. Once the tissue sunk in the 30% sucrose solution, it was placed in a mold, in optimal cutting temperature compound and allowed to freeze on dry ice. Cryosections (10-μm thick) were then cut and adhered to glass slides. Cryosections were permeabilized and stained for 30 min at room temperature in 0.5% Triton X-100 in PBS with Alexa Fluor-568 phalloidin or Alexa Fluor-647 phalloidin diluted 1:1000, and then blocked in 10% NGS in PBS overnight at 4℃. Tissue was incubated with primary antibody against NMIIA (rabbit), AQP5 (rabbit), or ZO-1 (rat) diluted 1:200-400 in 10%NGS for 4 h at room temperature, rinsed with PBS three times, stained with goat anti-rabbit or goat anti-rat secondary antibodies conjugated with Alexa Fluor-488 or Alexa Fluor-568 diluted 1:1000 in 10%NGS for 1 h. Tissue was then rinsed with PBS three times and mounted using mounting media (Fluoromount-GTM with DAPI) on a glass slide with no. 1.5 coverslips.

Microscopy was performed using a Nikon ECLIPSE Ti2 inverted fluorescence microscope (Tokyo, Japan), outfitted with a spinning disk confocal scan head (CSU-W1; Yokogawa Electric; Tokyo, Japan), 40x 1.30 N.A., 60x 1.40 N.A., and 100x 1.49 N.A. objective lens (Nikon), and ORCA-Fusion BT Digital CMOS camera (Hamamatsu Photonics; Hamamatsu, Japan). NIS-Elements imaging software (Nikon) was utilized for image acquisition and analysis.

### 7. Western blot analysis

The lacrimal and parotid glands were isolated under the anesthesia as described above. These glands were homogenized in ice-cold RIPA lysis buffer (25 mM Tris HCl pH 7.6, 150 mM NaCl, 1% NP-40, 1% sodium deoxycholate, 0.1% sodium dodecyl sulfate (SDS)) containing protease inhibitors (AEBSF, aprotinin, bestatin, E-64, leupeptin, and pepstatin A) and 1 mM phenylmethylsulfonyl fluoride; the homogenates were then incubated on ice for 15 min. The homogenates were spun at 14,000 g for 10 min. Supernatants were collected, and the protein concentrations were determined by the method of Bradford (52). The supernatants were used for Western blotting. Twenty-μg protein samples were separated by SDS-PAGE using a Mini-Protean 3 Cell system (Bio-Rad Laboratories, Inc.; Hercules, California, USA). After electrophoresis, the separated proteins were transferred onto a nitrocellulose membrane. The blots were blocked at room temperature for 1 h in 5% skim milk and then probed with a primary antibody, anti-NMIIC (rabbit; diluted 1:250), anti-AQP5 (rabbit; diluted 1:2,000), anti-ZO-1 (rabbit; diluted 1:500), or α-tubulin (mouse; diluted 1:1,000), overnight at 4℃. The blots were washed three times with Tris buffered saline (pH 7.4) containing 0.1% Tween 20, probed at room temperature for 1 h with anti-rabbit IgG antibody conjugated with Alexa Fluor^®^ 790 (diluted 1:50,000) for NMIIC, ZO-1 and AQP5, or anti-mouse IgG antibody conjugated with Alexa Fluor^®^ 680 (diluted 1:50,000) for α-tubulin, and washed again. Images were acquired using Odyssey CLx Imager (LI-COR Biosciences; Lincoln, NE, USA). The intensity of bands was measured with Image Studio version 5.2 (LI-COR Biosciences).

### 8. Administration of 4-HAP

An NMIIC activator, 4-HAP, was dissolved in ethanol and then diluted 100-fold with PBS. 4-HAP was injected intraperitoneally (i.p.) at a dosage of 1 mg/kg (volume: 0.05 mL/10g body weight) for 7 consecutive days based on a previous paper (22). Tear volume measurement and isolation of the lacrimal glands were performed on the last day of injection. The vehicle group was administered the same volume of PBS containing 1.0% ethanol.

### 9. Statistical analysis

Data are presented as the mean ± standard deviation (S.D.) (N = sample size). Statistical comparisons were made using a two-tailed Student’s *t-*test (Figure 2A-iii, 3B-iii, 4C-iii, 4F-ii, and 4F-iii); *t-*test with Welch’s correction (Figure 3A-iii); two-way ANOVA followed by Sidak’s multiple comparisons test (Figure 2A-ii, 3B-ii, and 4C-ii); or two-way ANOVA followed by Fisher’s LSD test (Figure 2D-ii and 2D-iii). Values (p) below 0.05 were regarded as statistically significant differences. These statistical analyses were performed using GraphPad Prism (GraphPad Software; La Jolla, CA, USA).

## Acknowledgments

The research was supported by JSPS Overseas Research Fellowship and Research Expenses for Overseas Trainees from Asahi University to Y.O., and the Center for Cell and Membrane Physiology, School of Medicine, at the University of Virginia through a start-up grant to S.E. We acknowledge Dr. Masanori Kashimata, Dr. Keitaro Satoh, Dr. Akiko Shitara, Dr. Haruna Nagase (Asahi University), and Dr. Bechara Kachar (National Institute of Deafness and Other Communication Disorders) for fruitful discussions.

## Author Contributions

Y.O. and S.E. conceptualized the study. Y.O. performed all the experiments with the help of G.H. D.R. created the quantification figure of fluorescence intensity in imaging. G.H. and D.R. contributed to maintaining NMIIC-GFP and NMIIC-KO mice. S.Z. assisted with the western blot preparation. Y.O. and S.E. wrote the manuscript, and made the figures for the experimental data, with contributions from all other co-authors.

## Competing Interest Statement

The authors declare that they have no competing financial interests.

## Supporting Information

### Supplementary Figure

**Supplementary Figure S1.**
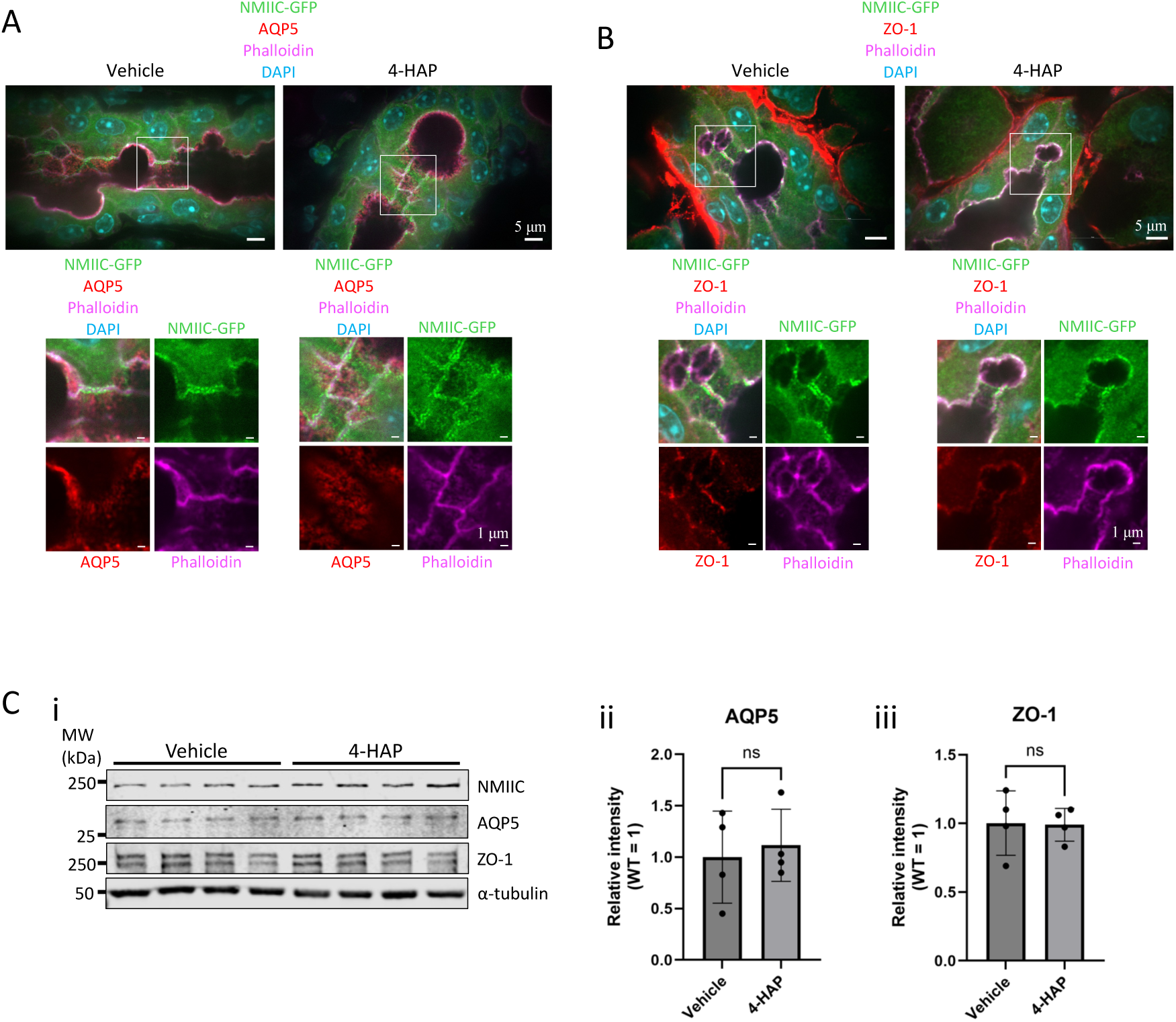
(A) Localization of AQP5 in lacrimal gland from 4-HAP- and vehicle-injected NMIIC-GFP mice. AQP5 (red) localizes on the apical membrane in ductal cells in the lacrimal gland from both groups. (B) Localization of ZO-1 in lacrimal gland from 4-HAP- and vehicle-injected NMIIC-GFP mice. ZO-1 (red) localizes at the apical junction of ductal cells in both groups. Phalloidin (magenta) represents F-actin. DAPI (blue) represents the nucleus. Scale bars, 5 μm in the upper image and 1 μm in the lower images. (C-i) Western blot of NMIIC, AQP5, ZO-1, and α-tubulin in lacrimal gland from 4-HAP- and vehicle-injected mice without CCh stimulation. 4-HAP-injected mice showed no NMIIC protein band and similar expression level of AQP5 and ZO-1 to vehicle-injected mice. (C-ii) Quantification of AQP5 band intensity. (C-iii) Quantification of ZO-1 band intensity. Student’s *t*-test. N = 4.

## Notes

### Competing Interest Statement

The authors have declared no competing interest.

